# A model for the propagation of seizure activity in normal brain tissue

**DOI:** 10.1101/2022.02.21.481321

**Authors:** Damien Depannemaecker, Mallory Carlu, Jules Bouté, Alain Destexhe

## Abstract

Epilepsies are characterized by paroxysmal electrophysiological events and seizures, which can propagate across the brain. One of the main unsolved questions in epilepsy is how epileptic activity can invade normal tissue and thus propagate across the brain. To investigate this question, we propose three computational models at the neural network scale to study the underlying dynamics of seizure propagation, understand which specific features play a role, and relate them to clinical or experimental observations. We consider both the internal connectivity structure between neurons and the input properties in our characterization. We show that a paroxysmal input is sometimes controlled by the network while in other instances, it can entrain the network activity to itself produce paroxysmal activity, and thus will further propagate to efferent networks. We further show how the details of the network architecture are essential to determine this switch to a seizure-like regime. We investigated the nature of the instability involved and in particular found a central role for the inhibitory connectivity.

We propose a probabilistic approach to the propagative/non-propagative scenarios, which may serve as a guide to control the seizure by using appropriate stimuli.

**Significance:** Our computational study shows the specific role that the inhibitory population can have and the possible dynamics regarding the propagation of seizure-like behavior in three different neuronal networks. We find that both structural and dynamical aspects are important to determine whether seizure activity invades the network.

We show the existence of a specific time window favorable to the reversal of the seizure propagation by appropriate stimuli.

## 1 Introduction

Epilepsy is one of the most common neurological disorder (Beghi, 2019), which can take numerous forms. It is associated with the presence of paroxysmal electrophysiological events and seizures, usually recorded in humans using the electroencephalogram (EEG). However, EEG recordings do not allow us to probe the activity of single neurons within the network. More recently, the recording carried out with microelectrode arrays made it possible to obtain spike information of the order of a hundred neurons (Peyrache et al., 2012; Dehghani et al., 2016).

[Il faudrait aussi citer le papier recent (Nat Neurosci) du groupe de Cash sur l’utilisation des neuro-pixels probes, qui permet d’avoir plusieurs centaines de neurones]

Such microelectrode recordings showed that neuronal activity during seizures does not necessarily correspond to synchronized spikes over the whole neuron population, as previously modeled (*Computational Neuroscience in Epilepsy*, 2008), including models at different scales from cellular to whole-brain levels (Depannemaecker, Destexhe, Jirsa, & Bernard, 2021; Depannemaecker et al., 2020). In fact, it turns out that the dynamics of neural networks during seizures are more complex (Jiruska et al., 2013), and still poorly understood. In particular, it is not known how the paroxysmal activity of the seizure does propagate, entraining other networks into seizure activity.

Here, we investigate this problem using computational models. We take as a starting point examples of seizures where the inhibitory network is strongly recruited, while excitatory cells’ firing is diminished. Fig.1 shows three seizures from a patient which were recorded using Utah-arrays, before resection surgery in a case of untractable epilepsy. From these intracranial recordings, 92 neurons have been identified and isolated and were classified into two groups: Fast-Spiking (FS) neurons and Regular-Spiking (RS) neurons, based on spike shape, autocorrelograms, and firing rates (Peyrache et al., 2012). Remarkably, direct cell-to-cell functional interactions were observed, which demonstrated that some of the FS cells are inhibitory while some of the RS cells are excitatory (see details in (Peyrache et al., 2012)). The three seizures shown in Fig.1 were taken from the analysis of (Dehghani et al., 2016) (see this paper for details), and are shown with the firing rate of each population of cells. During the seizure, we can observe a plateau of high activity of FS cells, and a strongly reduced activity of RS cells. This phenomenon of unbalanced dynamics between RS and FS cells was only seen during seizures in this patient (Dehghani et al., 2016). It shows that, in these three examples, the seizure was manifested by a strong “control” by the inhibitory FS cells, which almost silenced excitatory RS cells. It is interesting to see that a very different conclusion would be reached if no discrimination between RS and FS cells was performed, which underlies the importance of discriminating RS and FS cells for a correct interpretation of the dynamics during seizures. Based on such measurements, we built a computational models based on larger number of cells in order to consider network effect that are not directly accessible with the recordings. We were interested in how seizure activity propagates or not, and what are the determinants of such propagation.

**Figure 1:**
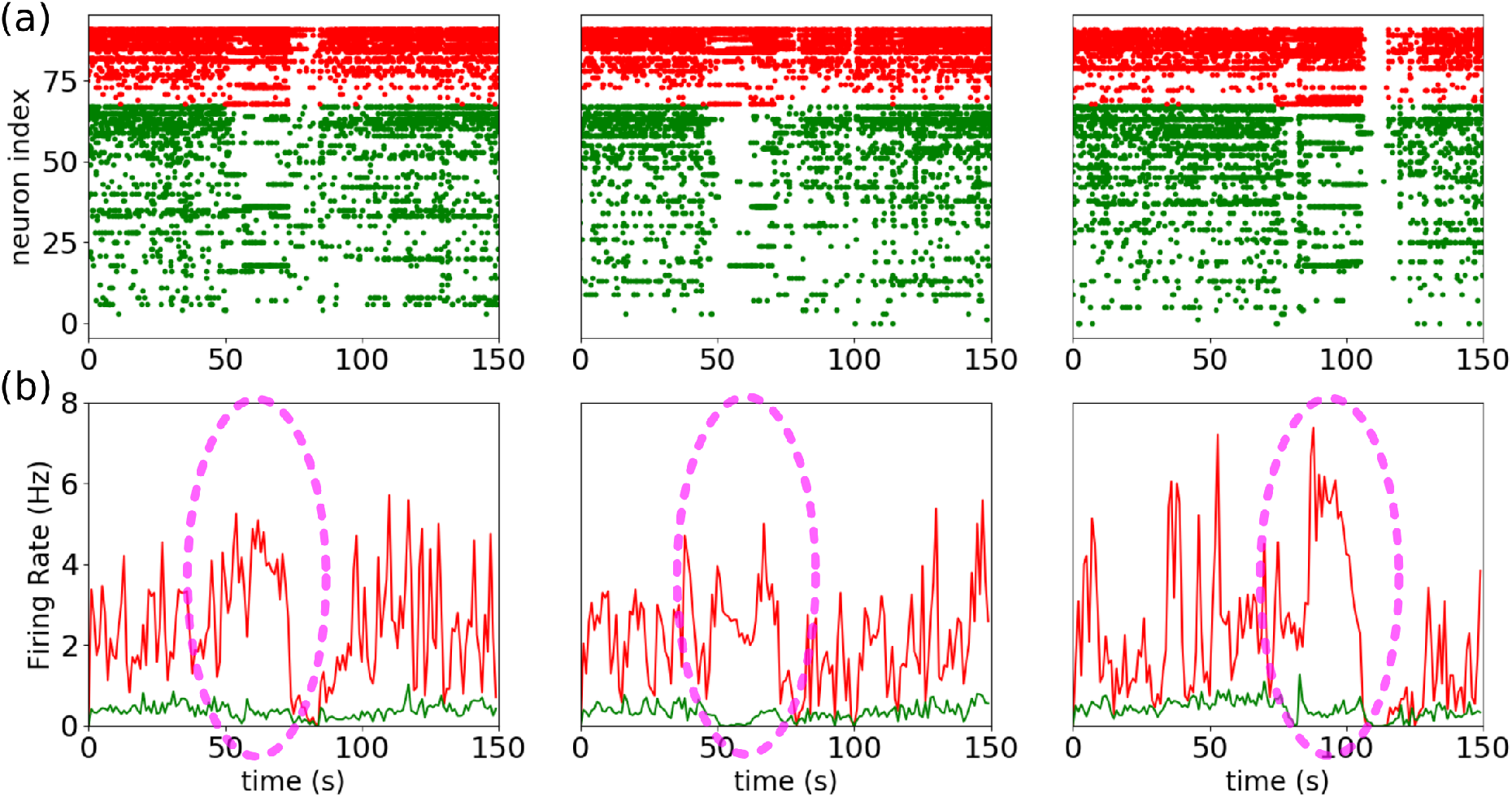
Examples of inhibitory recruitment during seizures: (a) Raster plot of three different seizures from the same patient, 92 neurones where identified 24 putatives inhibitory cells (red) and (68) putative excitatory cells (green) (b) Corresponding firing rate of the putative inhibitory population (red) and the putative excitatory population (green). A plateau of high activity of the putative inhibitory cells can be observed during the seizure (highlighted in dashed purple oval). Done with data from (Dehghani et al., 2016).

The region of the brain where the seizure starts is called the seizure focus, although in certain patients it is distributed over several foci (Nadler & Spencer, 2014), then the seizure spreads to other regions of the brain. When another such region is reached, it can in turn be entrained into seizure activity, in which case the seizure activity propagates. It can also control it (as in Fig.1), in which case the seizure would remain confined to a more restricted brain region.

In order to gain understanding of the dynamics underlying these two scenarios, we study the response of networks using three different neuron models (Adaptive exponential Integrate and fire (AdEx), Conductance-based Adaptive Exponential integrate-and-fire (CAdEx), and Hodgkin-Huxley (HH) models), interacting through conductance-based synapses, to an incoming paroxysmal (seizure-like) perturbation. We observe two types of behavior that we represent in Fig.2: one where the incoming perturbation successfully entrains the excitatory population, thus making its activity stronger than the input, and the other where only the inhibitory population strongly increases its activity, thus controlling the perturbation. In the first case, where the excitatory population discharges very strongly, it is therefore likely to transmit, or even amplify the perturbation transmitted to the next cortical column We have therefore called this situation the propagative scenario. In the opposite case, where the firing rate of the excitatory population remains much lower than the perturbation, the seizure-like event will not spread to the neighboring region, we therefore call this situation the non-propagative scenario. We then propose a more precise approach, based on the AdEx network, that mixes structural and dynamical ingredients in order to unravel key aspects of the mechanisms into play. Focusing on the different input connectivity profiles for each node in the network, we are able to build separate groups of neurons that display significantly different dynamics with respect to the perturbation. Finally, we study the possibility of a proactive approach, based on the application of an extra stimulus with the aim of reversing the propagative behavior, thus controlling the spread of the seizure.

**Figure 2:**
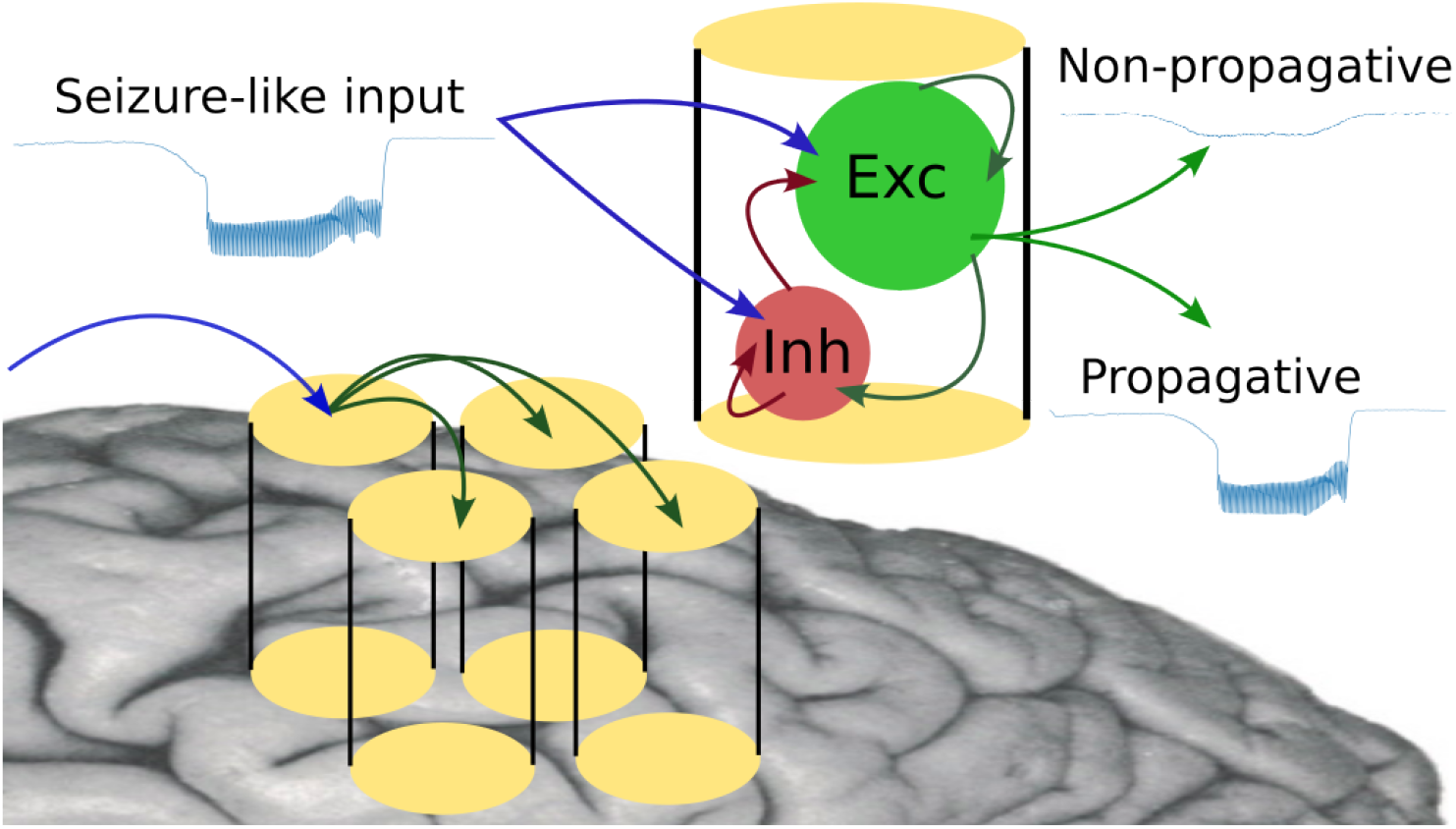
Cartoon of the modeled scenarios.

## 2 Material and methods

### 2.1 Computational models

We use for this study a mathematical model of electrophysiological activity based on ordinary differential equations, describing the dynamics of the neurons’ membrane potential through their interactions.

Each neuron model in the network is described by Eq.(1) and Eq.(2), the Adaptative Exponential integrate and fire (AdEx) model (Brette & Gerstner, 2005; Naud, Marcille, Clopath, & Gerstner, 2008).

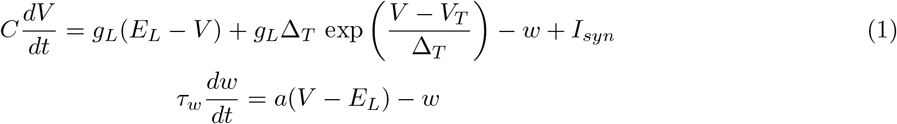

When the membrane potential crosses a threshold, a spike is emitted, and the system is reset:

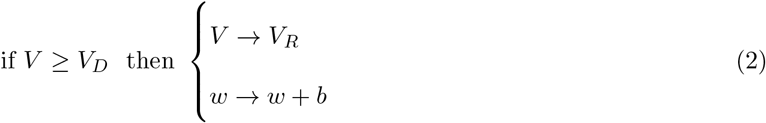

Parameters used for the excitatory (RS) and inhibitory (FS) populations are respectively *V*_*t*_ = −50 mV and *V*_*t*_ = −48 mV, *D*_*t*_ = 2 mV and *D*_*t*_ = 0.5 mV, *b* = 100 pA and *b* = 0 pA, and *τ*_*w*_ = 1000 ms for RS. For both population: *C*_*m*_ = 200 pF, *g*_*l*_ = 10 nS, *E*_*l*_ = −65 mV, *a* = 0 nS, *V*_*reset*_ = −65 mV, *t*_*refractory*_ = 5 ms.

In order to compare some of the results obtained with the AdEx model we used two other models of neuronal activity. First the Conductance-based Adaptive Exponential integrate-and-fire model (CAdEx), which solves some of the limitation of the AdEx model (Górski, Depannemaecker, & Destexhe, 2021). The equations read:

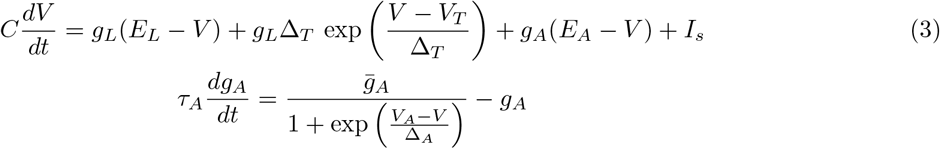

When the membrane potential crosses a threshold, a spike is emitted, and the system is reset as in:

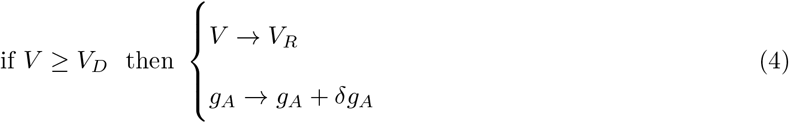

Parameters used for inhibitory (FS) populations are: *g*_*l*_ = 10 nS, *E*_*l*_ = −65 mV, *V*_*T*_ = −50 mV, *ga* = 0. nS, *E*_*A*_ = −70 mV,, *δg*_*A*_ = 0 nS, *C* = 200 pF, Δ_*T*_ = 0.5 ms, *V*_*A*_ = −45 mV, *Is* = 0.0 nA, *refractory* = 5 ms, *V*_*reset*_ = −65 mV, *tau*_*A*_ = 0.01 ms, Δ_*A*_ = 0.5 mV and for the excitatory (RS): *g*_*l*_ = 10 nS, *E*_*l*_ = −65 mV, *V*_*T*_ = −50 mV, *δg*_*A*_ = 1 nS, *E*_*A*_ = −65 mV, *δg*_*A*_ = 1 nS, *C* = 200 pF, Δ_*T*_ = 2 mV, *V*_*A*_ = −30 mV, *Is* = 0.0 nA, *t*_*refractory*_ = 5 ms, *V*_*reset*_ = −65 mV, *tau*_*A*_ = 1.0 s, Δ_*A*_ = 1 mV Then we use the Hodgkin-Huxley model (Hodgkin & Huxley, 1952), hereafter denoted HH, with the following equations:

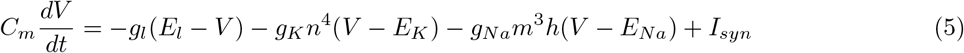

with gating variables (in ms):

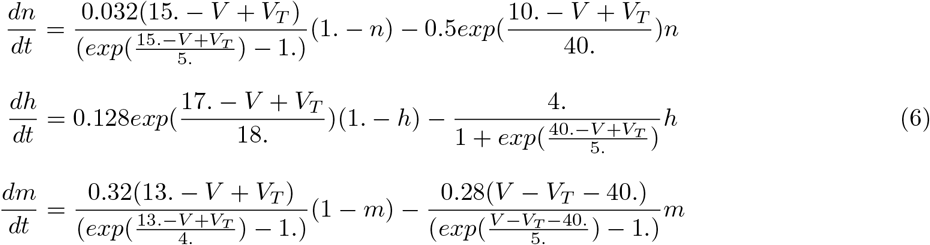

With *C*_*m*_ = 200 pF, *E*_*l*_ = −65 mV, *E*_*Na*_ = 60 mV, *E*_*K*_ = −90 mV, *g*_*l*_ = 10 nS, *g*_*Na*_ = 20 nS, *g*_*K*_ = 6 nS, *V*_*T exc*_ = −50 mV, *V*_*T inh*_ = −52 mV.

For all types of neuron models, the parameters have been chosen in biophysical range (see (Naud et al., 2008; Górski et al., 2021; Hodgkin & Huxley, 1952; Hille, 1992)) in order to keep the basal asynchronous irregular activities (Brunel, 2000) into a range of firing rate coherent with experimental observations (El Boustani, Pospischil, Rudolph-Lilith, & Destexhe, 2007; Destexhe, 2009; Zerlaut, Chemla, Chavane, & Destexhe, 2018).

The network is built according to a sparse and random (Erdos-Renyi type) architecture where a fixed probability of connection between each neurons is set to 5%. We consider a network model of ten thousand neurons, built according to specific properties of the cortex. This network is made of an inhibitory (FS) and an excitatory (RS) population, respectively representing 20% and 80% of the total size of the system as previouly done in (Carlu et al., 2020) The communication between neurons occurs through conductance-based synapses. The synaptic current is described by:

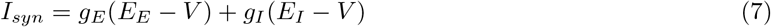

Where *E*_*E*_ = 0 mV is the reversal potential of excitatory synapses and *E*_*I*_ = −80 mV is the reversal potential of inhibitory synapses. *g*_*E*_ and *g*_*I*_, are respectively the excitatory and inhibitory conductances, which increase by quantity *Q*_*E*_ = 1.5 nS and *Q*_*I*_ = 5 nS for each incoming spike. The increment of conductance is followed by an exponential decrease according to:

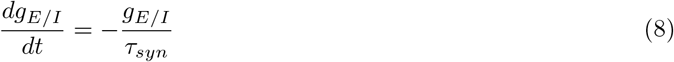

with *τ*_*syn*_ = 5 ms

The network thus formed receives an external input, based on the activity of a third population (excitatory) of the same size as the excitatory population. Each of its neurons is connected to the rest of the network according to the same rule as mentioned earlier (fixed probability of 5 % for each connection). This external population produces spikes with a Poissonian distribution at a given tunable rate. The external perturbation that mimics the incoming seizure occurs through the augmentation of this firing rate.

The shape of the latter is described by:

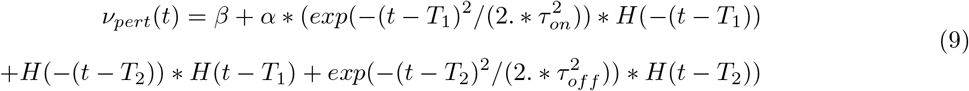

where *H* is the Heaviside function and *β* = 6 Hz is the basal constant input. This function takes the general form of a high plateau, where *T*_1_ and *T*_2_ are the times when the perturbation reaches its beginning and end respectively, and *α* defines its maximal height. *τ*_*on*_ and *τ*_*off*_ are respectively time constants associated with the exponential rise and decay of the perturbation.

For all 3 types of networks, it is possible to have different connectivities (i.e different set of random connectivities) and realizations of Poisson drive (i.e the generator of the Poisson noise can vary). It is also possible to fix the seed of the noise either for the connectivity or for the Poisson drive (or both) to analyse specific conditions.

We create network connectivities by allowing a 5% chance of connection between any 2 neurons, which will indeed lead to an average of 5% of connection, but with some variation. Some neurons can have more afferent connections from inhibitory neurons than others, which will make them more inhibited, and the same goes with excitatory connections, creating a variation between neurons due to the random nature of the network.

### 2.2 Coarse graining and continuous analysis

In order to analyse in details what mechanisms are at play in the network during a seizure-like event, we resort to a combination of two methods : a so-called *structural coarse-graining*, that is we gather neuron models in *n* groups according to their inhibitory in-degree (the number of inhibitory connections they receive, as introduced before, and we study their time evolution through statistics of their membrane potential (mean and alignment) over these groups. In other words, at each integration time step, we will obtain *n* values of mean membrane potentials, one for each group, as well as *n* values of Kuramoto order parameter (measuring alignment in groups).

To obtain the Kuramoto order parameter, we first transform the single neurons membrane potentials into phase variables by applying a linear mapping 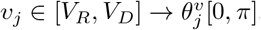. Then the Kuramoto order parameter is computed through the following equation:

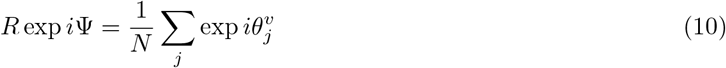

*R* ∈ [0, 1] gives the degree of “alignment” (if it persists in time, one would say synchronization) : *R* = 1 implies full alignment, while *R* = 0 implies no alignment whatsoever. Ψ ∈ [0, *π*] tells us the mean phase of the transformed variables (directly related to the mean membrane potential).

Let us mention one caveat here. The membrane potentials are not mapped on the full circle, to avoid artificial periodicity of the obtained angles: having *V* = *V*_*R*_ is not the same as having *V* = *V*_*D*_. One may thus ask why such a measure is used instead of the usual measures of dispersion such as the standard deviation. We use the Kuramoto order parameter because it gives a naturally normalized quantity, thus allowing a direct comparison of what is happening at each time step.

### 2.3 Code Accessibility

The code/software described in the paper is freely available online at [URL redacted for double-blind review]. The code is available as Extended Data and is run on Linux operating system.

## 3 Results

We start by showing how, in networks of various neuron models, a paroxysmal external stimulation can trigger a seizure, depending on various parameters. We show how the situations differ from model to model and what are the common features. Then, we propose a structural analysis based on mean firing rates of individual neuron models to guide a particular coarse-graining, which we use as a filter to observe the dynamics and gain further understanding, from both qualitative and quantitative perspectives. Finally we show how this study can guide proactive approach to reduce the chances of crisis propagation.

### 3.1 Propagative and Non-propagative behaviors

Throughout this study, we assume that the networks depicted in the previous section represent a small cortical area receiving connections from an epileptic focus. Specifically, the arrival of the seizure is modeled by a sudden rise in the firing rate of the external (afferent) Poisson region where the crisis comes from, or originates. In other words, we are not concerned with how crises *originate* (epileptogenesis), but how they *can propagate*. Therefore, we will frame our analysis into two main scenarios: *propagative, i.e* the network develops an excitatory firing rate greater than the input, which makes it able to propagate the crisis to efferent regions, and *non-propagative* behavior where the excitatory firing rate is lower than the input, thus attenuating the incoming signal. As described in the method section, the perturbation starts with an exponential growth followed by a plateau and ends with an exponential decrease, going back to the basal level : see blue curves in Fig.3. We show in this figure the response of the various networks to this type of perturbation.

**Figure 3:**
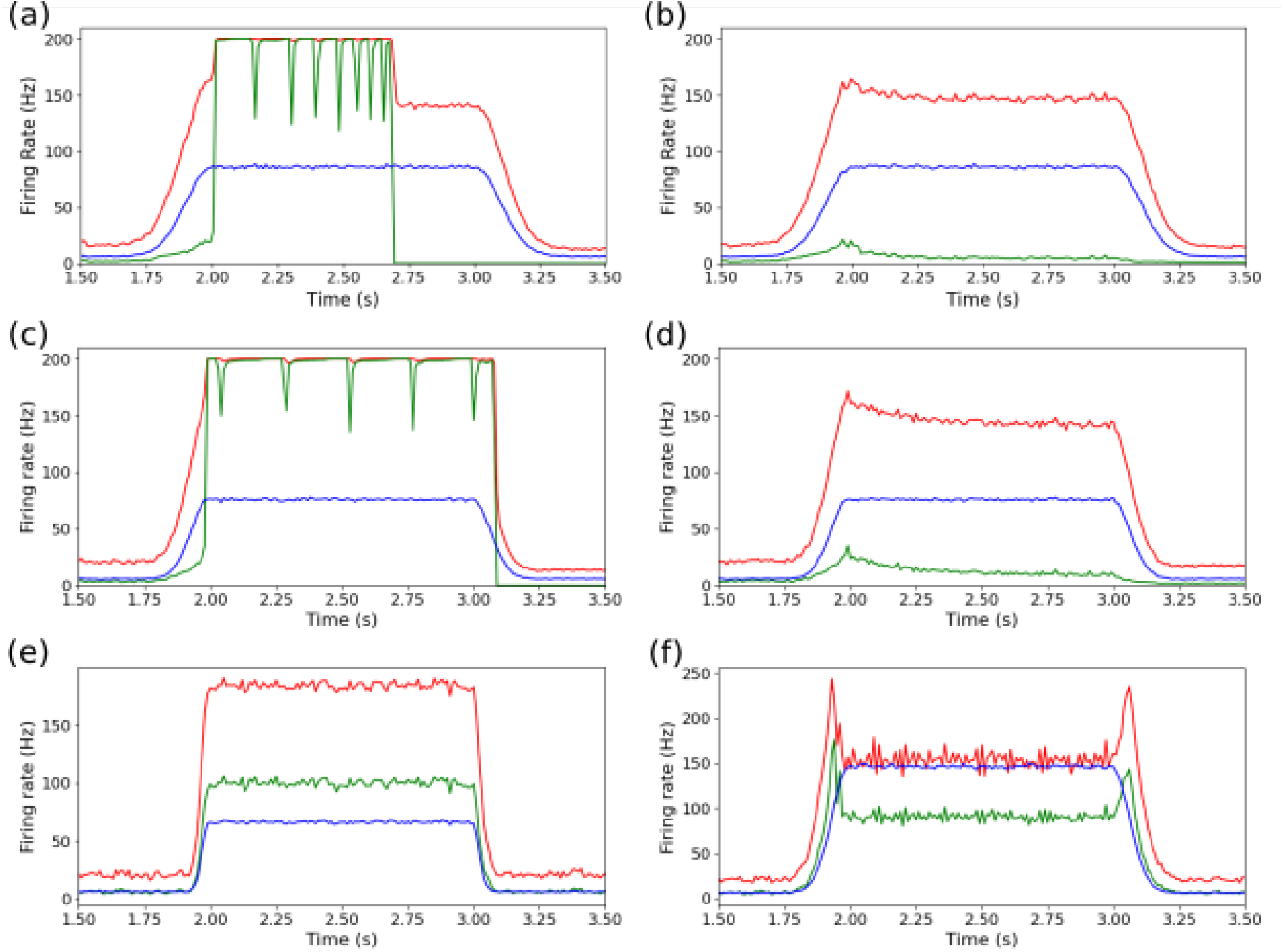
Firing rate of the network populations in response to a perturbation. (in blue the incoming perturbation, in green excitatory and in red inhibitory populations): propagative and non-propagative behaviors (respectively left and right columns) for AdEx model ((a) and (b)), with amplitude of perturbation *α* = 80*Hz* and *τ*_*on/off*_ = 100*ms*; CAdEx model ((c) and (d)) with *α* = 70*Hz* and *τ*_*on/off*_ = 80*ms*; HH model (e) with *α* = 60*Hz* and *τ*_*on/off*_ = 60*ms* and (f) with *α* = 140*Hz* and *τ*_*on/off*_ = 60*ms*. [FIGURE MODIFIED]

Here we can distinguish between two classes of macroscopic differences between propagative and non-propagative scenarios.

In the first class (for AdEx and CAdEx) the difference is binary. That means the network either features a very strong increase in the firing rate of the inhibitory *and* excitatory populations, or the sharp increase in the firing rate only concerns the inhibitory population, thus strongly limiting the activity of the excitatory population (consequently preventing the seizure from spreading to the next region). From this perspective, the propagative scenario can be understood as a loss of balance between excitatory and inhibitory firing rates, which the network struggles to find once the excitatory population has reach very high firing rate. Interestingly these two scenarios can occur for the same global shape of the perturbation but changing only the noise and network realizations. It must be noted that the 200*Hz* maximum frequencies measured here are the results of the temporal binning of the global spiking dynamics, taken as *T* = 5*ms*, which corresponds to the refractory time of the single neurons in Adex and CAdEx. Upon choosing a shorter binning, *e.g T* = 1*ms*, higher frequency peaks are observed, going up to 800*Hz*, thus hinting at overall faster dynamics. In the second class (HH) there is a rather continuous difference between propagative and non-propagative behaviors as can be seen in Fig.4(c), depending on the amplitude of the perturbation.

**Figure 4:**
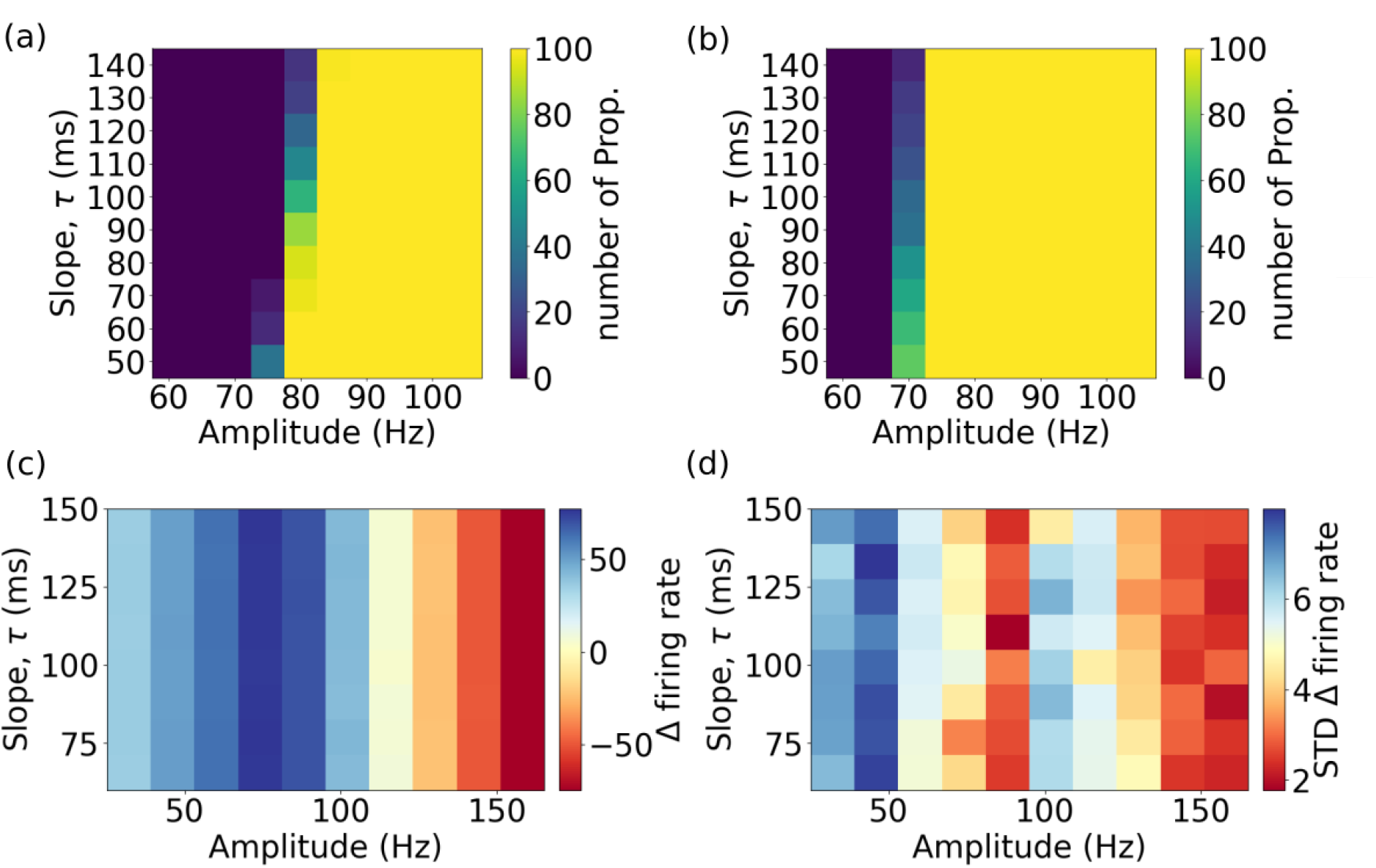
Grid search on the amplitude and slope of the incoming perturbation for each network. Panels (a) and (b) show the percent of realizations which propagate, respectively for Adex and CAdEx networks. Panels (c) and (d) show respectively, for HH networks, the means and standard deviations (over realizations) of the difference of firing rates between excitatory and Poisson populations (Δ *firing rate* = *ν*_*e*_ − *ν*_*P ois*_), averaged over the length of the plateau.[FIGURE MODIFIED]

### 3.2 Influence of the perturbation’s shape

To study how the shape of the perturbation affects the networks response, we screened in Fig.4 different time constants of the exponential growth rates and maximum amplitude of the plateau with 100 seeds (for both network and noise realizations for each couple of values, and probed, in the case of AdEx and CAdEx (respectively (a) and (b)), the number of realizations which yield propagative behavior, as it shows binary possible scenarios. Meanwhile, in the HH case, the perspective is a little different : we chose to show two figures, displaying means and standard deviations over realizations of the difference in firing rate between excitatory and Poisson populations (averaged over the plateau), Δ *firing rate* = *ν*_*e*_ − *ν*_*P ois*_ (respectively (c) and (d)). As can be expected, for all networks (AdEx, CAdEx and HH) the amplitude of the perturbation plays a determinant role in the type of scenario we eventually find (propagative or not), however in opposite directions and of different nature. Indeed, for both AdEx and CAdEx, increasing the amplitude increases the chance of having a propagative scenario for a fixed slope, in a binary fashion, while in the case of HH the contrary is observed, and in a continuous fashion. Also, we observe a slight coupling effect between slope and amplitude : for higher amplitudes, the propagation range extends to slower perturbations. On the contrary, in the HH network, it seems that the slope does not play any major role, hinting at a much less dynamical effect : the difference manifest themselves as local equilibria of the networks under considerations, reached no matter the time course. Moreover, the standard deviations, besides showing no clear dependence on neither amplitude nor slope, are very small compared to the means, thus evidencing that noise neither plays any significant part here. These observations highlight once again deep differences between the two types of network and their respective phenomenology.

Interestingly, in the case of AdEx and CAdEx, there exists a limit, bi-stable region here, around 80*Hz*, where the perturbation may or may not propagate in the network, depending on the noise realisation. Thus, the scenario does not trivially depend on the amplitude and time constants of the perturbation in this region, which makes the latter a perfect test bed to study more deeply the internal mechanisms at play, and will thus be the main focus of the remainder of this paper.

### 3.3 Influence of structural aspects on the dynamics

In the following, we turn our attention to the bi-stable region of AdEx networks, where the two scenarios are present, and investigate what can be the source of the divergence. There are two main differences between the simulations under consideration: the realization of the network structure and the realization of the external input, as both rely on random number generators. We have therefore successively fixed each of them, and observed that the two behaviors were still present. Also, the global scenarios were indistinguishable from those showed so far. First, this allows us to fix the network structure (which will become determinant in this part) without losing the richness of the phenomenology. Second, this tells us that what shapes the distinction between the two phenomena is more complex than a single question of structure, or realization of the input. Another perspective is then needed to explore the internal dynamics of the network in both scenarios. As the models into consideration have very large number of dimensions, as well as quite intricate structures, brute force analytical approaches are simply not conceivable.

Let us then take a step back and investigate the relationship between the firing rate of each neuron and its number of afferent (input) connections for the three kinds of input: excitatory 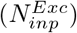, inhibitory 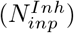 and Poissonian 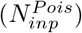. Fig.5(a) shows the average firing rates 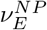 and 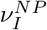 measured over the whole non-propagative (NP) scenario for each neuron in the AdEx network (simply defined as the total number of spikes divided by the total integration time, after having discarded a transient), plotted as a function of the three different connectivities.

**Figure 5:**
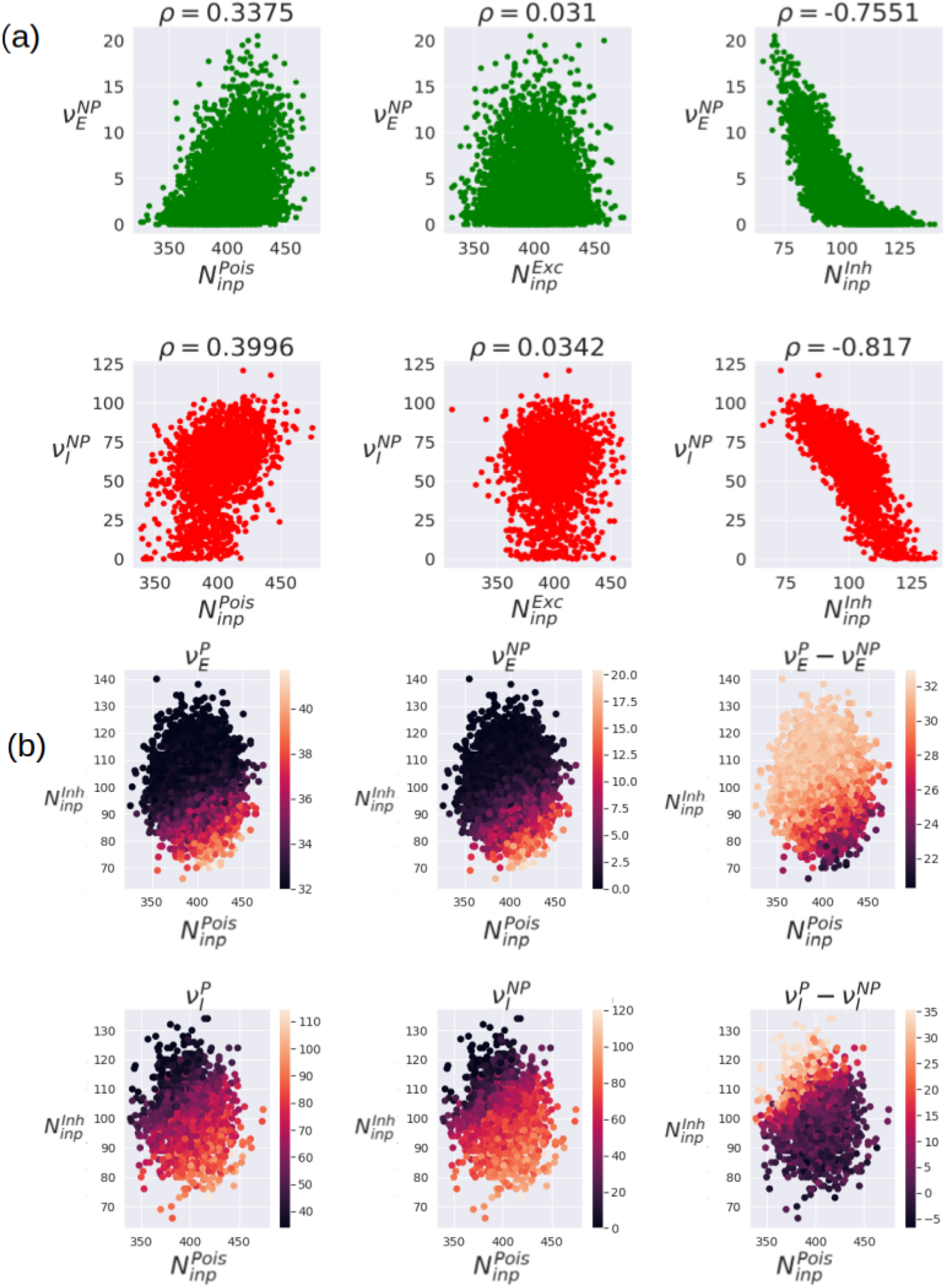
Influence of connectivity on single neurons firing rates: (a) Influence of poissonian 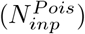, excitatory 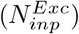 and inhibitory 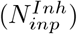 in-degree on the firing rates of excitatory neurons 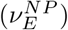, and inhibitory ones 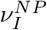 in non-propagative scenario (NP) of the AdEx network. The standard pearson correlation coefficient *ρ* is estimated. (b) Time averaged single neuron firing rates and differences in propagative vs non-propagative regimes, as a function of both inhibitory and poissonian in-degrees.[FIGURE MODIFIED]

Note here that averaging over simulations for the sake of robustness might be a delicate matter, as we might lose constitutive differences in the process. As we are dealing with highly variable situations, we have to make compromises between generalizability and relevance. Therefore, we start with a single realization to then guide larger and more systematic investigations.

Interestingly, we see a much stronger influence coming from the inhibitory in-degree than from the Poissonian and excitatory ones. Counter-intuitively, it even seems that excitatory in-degree has barely any effect at all on total measured firing rates. Indeed, from the point of view of Pearson’s correlation, inhibitory in-degree is much more (anti)-correlated with the firing rate than the excitatory in-degree (almost no correlation) or the Poissonian in-degree (little correlation). Note that we observe the same structure for propagative situations (results not shown). Based on these results, we can analyze whether the most salient in-degrees (inhibitory and Poissonian) has any influence on the *difference* between propagative and non-propagative situations, see Fig.5(b). Here, we see that the global dependency of the average single neuron firing rates on inhibitory and Poissonian connectivity does not qualitatively change between propagative and non-propagative regimes. However, the differences *ν*^*P*^ − *ν*^*NP*^ display an inverse dependency on both variables: despite maintaining a qualitative similarity between first and second columns, the crisis tends to compensate the initial disparity of firing rates. In other words, the neurons that are initially less firing, due to their structural properties, are the most impacted by the crisis. Furthermore, it must be noted that, although there is no correlation between inhibitory and Poissonian in-degrees (as can be expected from random connectivities), the third column highlights that they both play a role in the single neurons long term dynamics.

To understand the effect of the inhibitory connectivity, we choose two points from Fig.4(a), one known to be always non-propagative, with *τ* = 70*ms* and *α* = 70*Hz*, the other to be always propagative, with *τ* = 70*ms* and *α* = 95*Hz*. In both situation, we varied the probability of connection from the inhibitory population between *p* = 0.04 and *p* = 0.06, results are shown in Fig.6. Note that the influence of incoming inhibitory connectivity shifts the boundary between propagative and non-propagative behaviors. This is an important influencing factor in relation to the shape of the perturbation and in particular its amplitude.

**Figure 6:**
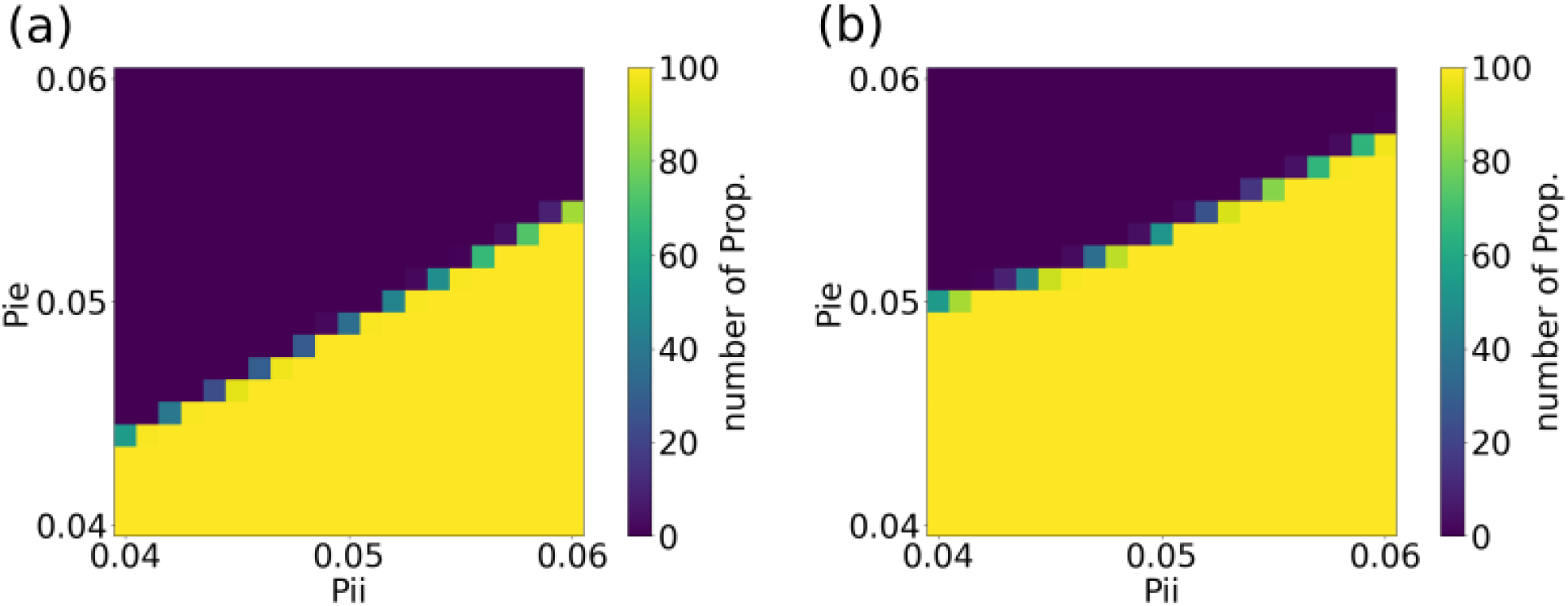
Grid search on the in-degree inhibitory probability of connection for the AdEx network. Percent of propagation with parameters: (a) *α* = 70*Hz* and *τ* = 70*ms*, (b) *α* = 95*Hz* and *τ* = 70*ms*, where for both figures Pie is the probability of connection from inhibitory to excitatory neurons and Pii is the probability of connection from inhibitory neurons to inhibitory neurons. [NEW FIGURE]

It is worth pointing that these results establish a clear link between structure and dynamics, but structure is by itself not a sufficient criterion to understand the underlying mechanisms. We threfore focus on the temporal evolution of the propagating activity.

Beforehand, we take a step back and probe whether the differences in the individual mean firing rates are associated with specific roles in the dynamics. To achieve so, we start classifying, for the AdEx network, the neurons’ indices in the raster plot according to the total number of spikes they emit during the whole simulation. We chose for this purpose a representative propagative scenario. The sequence of propagation of the perturbation then appears visually in Fig.7(a). We observe, in the case of propagation, a fast cascade (consistent with the experimental observations (Neumann et al., 2017)) : these simulations show that some neurons are quickly “entrained” in a sequence at the onset of the seizure.In addition, there is no perfect synchronization of the action potentials of all neurons. This is an interesting result, coherent with experimental observations on epilepsy (Jiruska et al., 2013).

**Figure 7:**
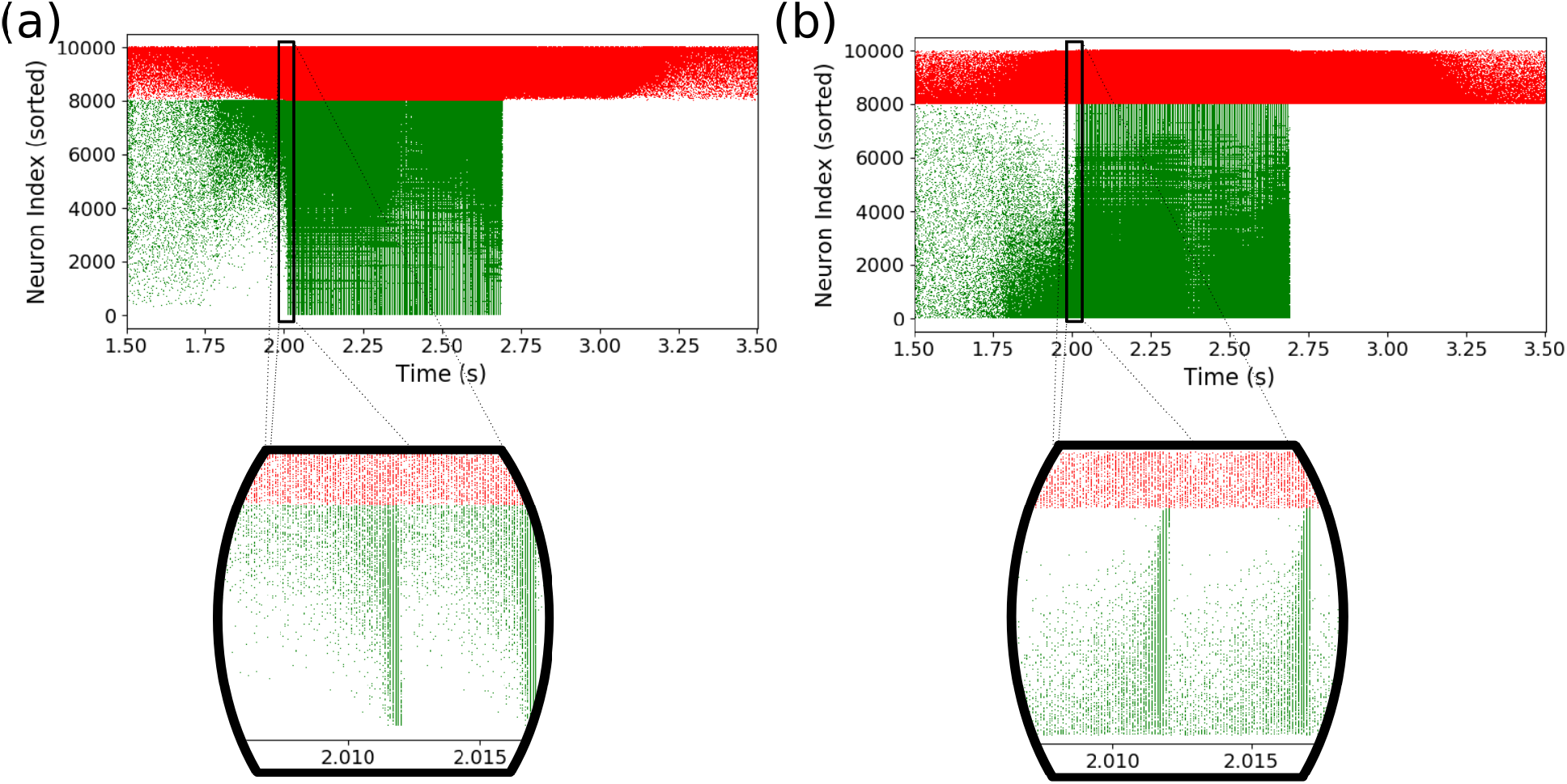
Dynamics in propagative situation (AdEx): (a) Raster plot of a simulation with propagative behavior, neuron indices are sorted according to the number of spikes during the simulation. A “cascade” phenomenon can be observed when zooming on the onset of the perturbation propagation in the excitatory population. (b) The same cascade phenomenon is observed when neuron indices are sorted in function of the number of inhibitory inputs received. Note that the absence of excitatory activity after the perturbation is due to a strong adaptation current (see Eq.(1), and Eq.(2)).

Secondly, we examine the same situation, but sorting neuron indices as a function of the number of inhibitory inputs they receive, as shown Fig.7(b), as it is the most influential structural feature we observed in our model. Here too, the cascade phenomenon is clearly visible, indicating that the inhibitory input connectivity has a central influence on the dynamics at play during the perturbation in the propagative scenario.

Fig.8 shows the same pictures for CAdEx and HH networks. We see here that CAdEx network’s behaviors are very similar to AdEx : sorting with firing rate or inhibitory in-degree gives very similar structures and we can distinguish here too the cascading effect at the onset of the perturbation, following the indices. HH networks show quite a different phenomenology. First the two sorting do not show the same structures, which hints at a more subtle mapping between inhibitory in-degree and long-term single neuron model dynamics. In the firing rate sorting, we can still distinguish 3 blocs of distinct activity, and thus of populations, corresponding to the 3 key periods of the simulation : before stimulation, at the onset, and during the stimulation. Interestingly it seems that before and during the stimulation different populations of neuron models are distinctly mobilized. While before the stimulation, the central neurons (with respect to their indices) are active, a double cascade contaminates the rest of them (towards higher and lower indices) at the onset, ending in a general surge of activity. This must be contrasted with the in-degree sorting panel, where the cascade is more unidirectional, as the main activity slides from low connectivity indices (less connected) to the higher ones, until all neurons fire. This emphasizes the importance of the perspective chosen to analyse complex behavior : none of these perspectives alone completely explain the intricate interplay between structure, long term, and short term dynamics.

**Figure 8:**
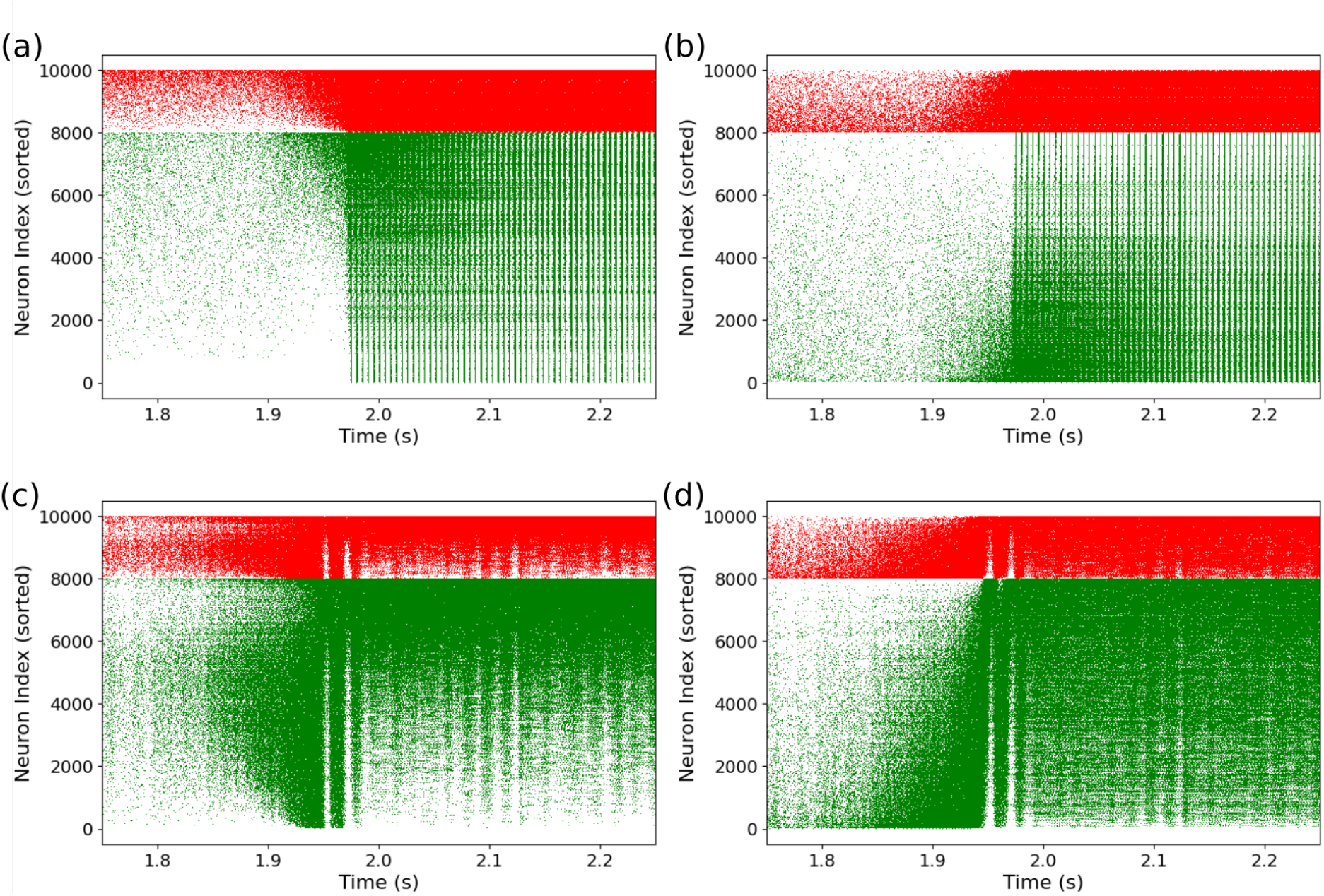
Dynamics in propagative situation (CAdEx and HH): Same plot as previously shown but for CAdEx network ((a) spike-sorting, (b) inhibitory in-degree sorting) and HH network ((c) spike-sorting, (d) inhibitory in-degree sorting). Cascade phenomena are still observable in panels (a),(b) and (d), hence showing its robustness, but not in (c), where propagation takes a slightly different form, highlighting the contrast induced by different perspectives on a single complex dynamics.[NEW FIGURE]

Altogether these results show the relevance of adopting a perspective based on the inhibitory in-degree : it gives an operational method to rank single neurons, and this ranking is clearly associated with specific dynamical features, hence allowing us to study the role of the internal organisation of the network before and during the paroxysmal event. As the cascade phenomenon is similarly visible in all types of networks, in the next section we focus on the AdEx network. We push further this analysis by comparing propagative and non-propagative scenarios, and make use of the continuous measures introduced in Material and Methods.

### 3.4 Continuous measures on subgroups of neurons

Focusing on the AdEx network, we first consider groups of neurons defined by their inhibitory in-degree. Note that these are somewhat artificial, as they are only statistical reflections of topological aspects of the network (i.e, there is no reason to think a priori that all neurons having *n* inhibitory inputs would have more privileged links among themselves than with those having a different numbe)r. However, they allow in principle a variable degree of categorization, based upon the sampling of the inhibitory in-degree distribution, which eventually leads to different levels of (nonlinear) coarse-graining (although we will consider only one such sampling here). Secondly, we switch our analysis to continuous variables, which allow a finer and more systematic analysis of the dynamics, as they don’t depend on spike times. Indeed, although spike timings are the most accessible collective measures in real-life systems, which make them the most fitted candidates for “transferable” studies, we want here to take advantage of the virtues of mathematical modeling to probe the insides of these simulations, to then be able to draw conclusions on more accessible observables. We focus here uniquely on membrane potentials, as they are the closest proxy of the firing dynamics in the network and chose to use two main measures based on them: the mean *µ*_*V*_ and a modified Kuramoto order parameter *R*, which gives a naturally normalized measure of instantaneous alignment (or similarity) of the membrane potentials. Both are defined in time, over a class of neurons. As randomness plays a crucial role in our simulations, through network structure as well as noise realization, it is important to control how much it affects the results we obtain. To achieve so, we start by fixing the network structure while averaging over noise realizations, and then average over connectivities while looking for noise realizations that lead to propagative and non-propagative scenarios for each structure.

#### Mean membrane potential in time

We show in Fig.9(a)-(b), the mean membrane potential *µ*_*V*_ defined for each group of excitatory (RS) and inhibitory (FS) neurons, in time.The top and bottom rows respectively refer to the averages and standard deviations over noise realizations (input), as the network structure is held fixed. (a) Propagative scenarios (b) non-propagative scenarios. All the data presented from now are obtained by regrouping neurons having the exact same inhibitory in-degree, thus corresponding to a discrete one-to-one sampling of the input distribution. Note that, given the network architecture under consideration, the number of afferent inhibitory synapses defined over both populations of neurons follows a Gaussian distribution with a mean around 100 connection. From that, we arrange neurons in groups of identical number of inhibitory connections, which gives us about 60 groups (varying with population and connectivities) containing at least 1 neuron.

**Figure 9:**
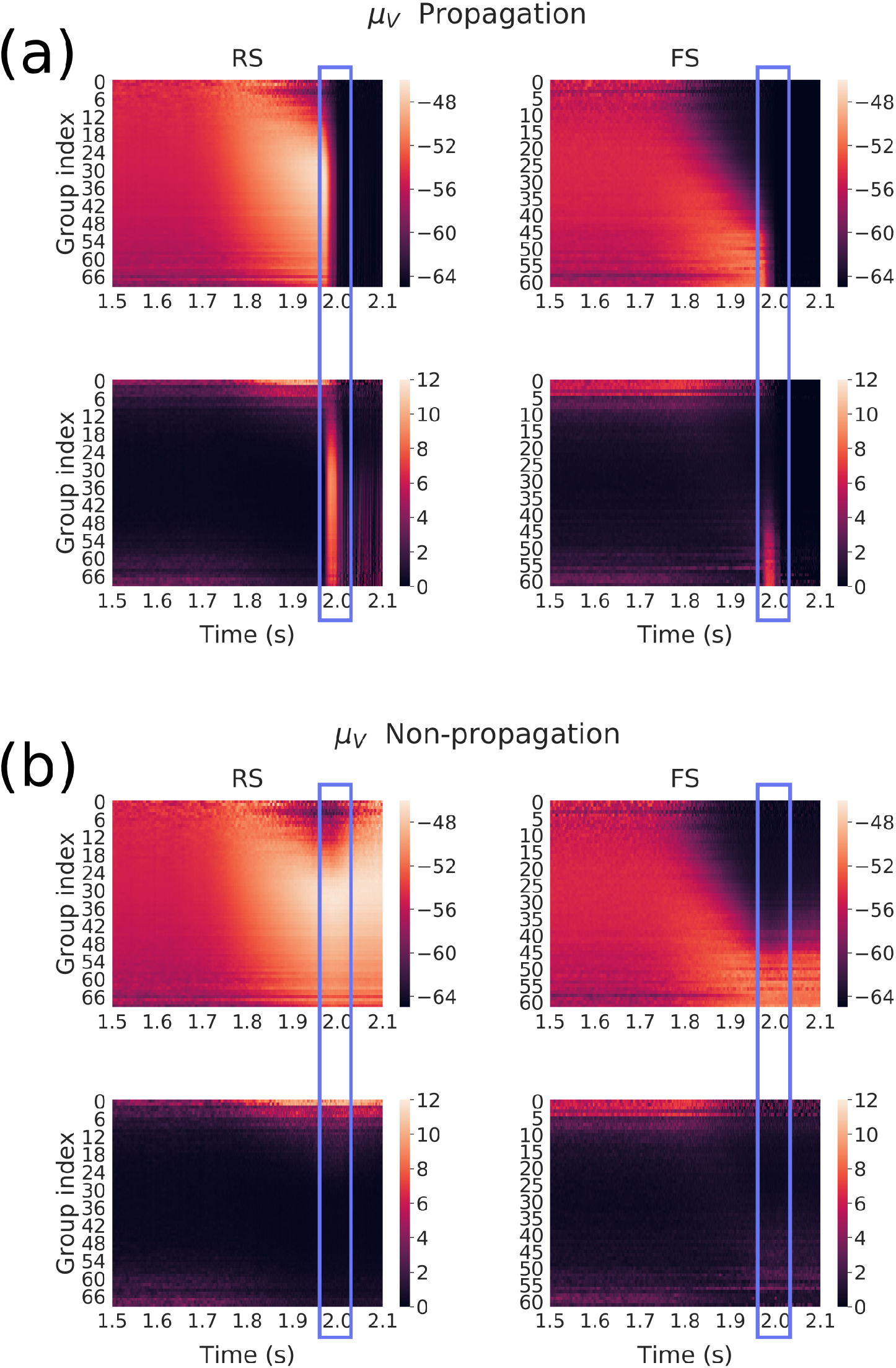
Mean membrane potential over subgroups of neurons (same network connectivity, different noise realizations) for each group defined as a function of their incoming inhibitory connections, averaged over 50 noise realizations (17 non-propagative and 33 propagative). Color maps correspond for each group to the average membrane potential (top) and standard-deviation (bottom) across noise realizations in the propagative situations (a) and non-propagative situations (b) for both excitatory (RS) and inhibitory (FS) populations. The blue rectangle highlight the (time) region where the system either switches to a propagative regime, or remains stable. [FIGURE MODIFIED]

To confirm that our results were not connectivity specific, we simulated 50 different networks with different connectivities (otherwise being identical) and found a couple of noise realizations for each corresponding to propagative and non-propagative scenarios. We applied the same method to create the different groups, but the number of said groups could differ due to the random variability in the connections. Therefore, many “extreme” groups are poorly represented among the various connectivities, which would make them hard to average over. We thus discarded them. The average and standard deviation of *µ*_*V*_ over the 50 different connectivities is shown in Fig.10(a)-(b). The white lines could be a weak manifestation of the previous effect, which made the standard deviation very high, coupled with the fixed range of color scales, imposed for the need of clarity. We observe that this figure looks very similar to Fig.9, which suggest that the results are not limited to a specific connectivity.

**Figure 10:**
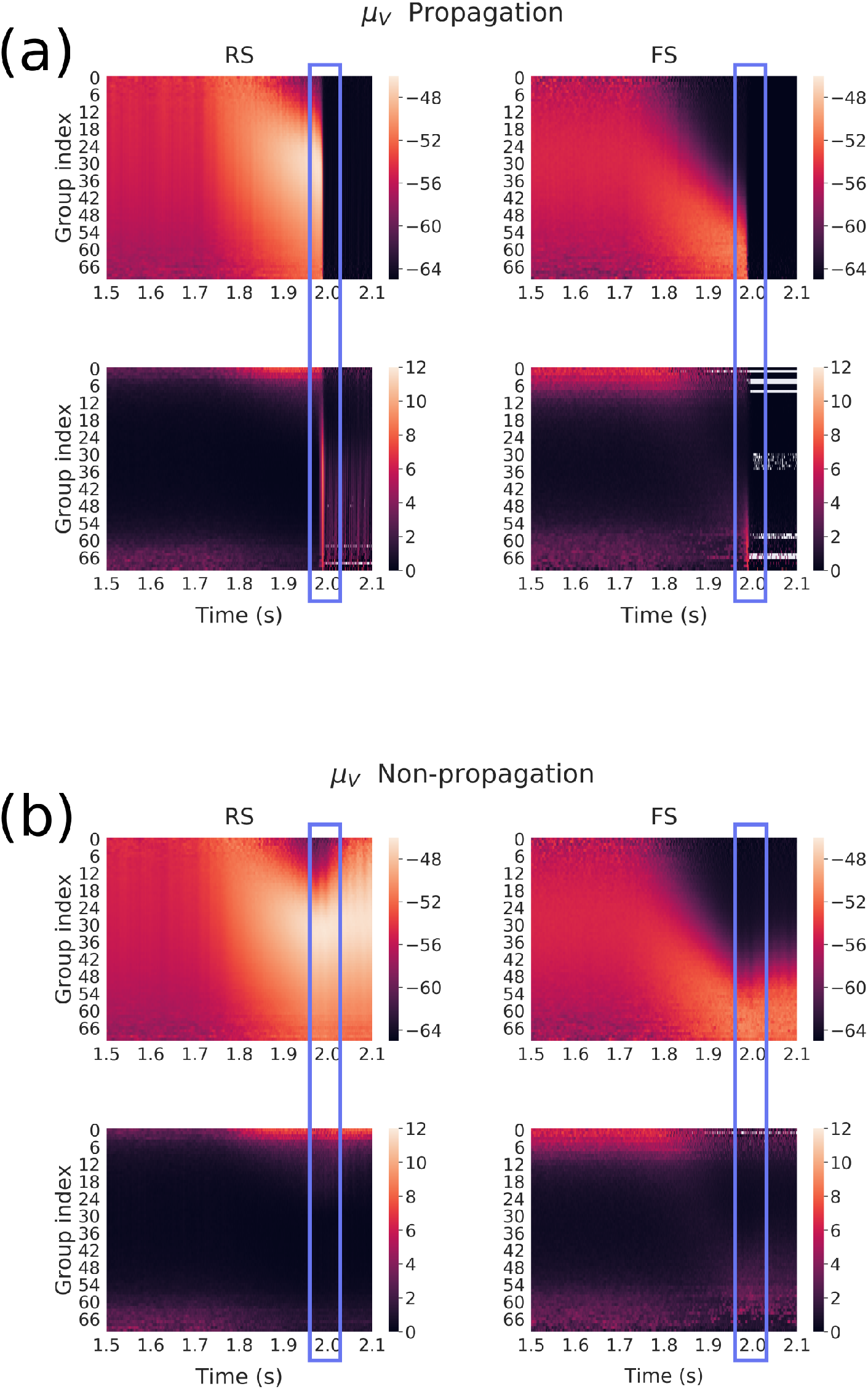
Mean membrane potential over subgroups of neurons (different network connectivities) for each group defined as a function of their incoming inhibitory connections. Here, we averaged over 50 network connectivities for which we found a couple of noise realizations corresponding to propagative and non-propagative scenarios. Color maps correspond for each group to the average membrane potential (top) and standard-deviation (bottom) across different connectivities in the propagative situations (a) and non-propagative situations (b) for both excitatory (RS) and inhibitory (FS) populations. The blue rectangle highlight the (time) region where the system either switches to a propagative regime, or remains stable. [NEW FIGURE]

We see from Fig.9 and Fig.10 that the inhibitory in-degree profile seems to play a major role in the overall dynamics. Indeed, as the perturbation is growing (starting 250*ms* before the maximum at 2*s*), we can first observe a strong increase of the mean membrane potential of all excitatory neurons, starting from low indices, then followed by a low-potential cascade, also starting from low indices and then contaminating to higher ones.

This latter effect is much clearer in the case of inhibitory neurons, where the cascade follows very well the input profile, in both propagative and non-propagative scenarios. Note that the low-potential area can be easily understood as a high-firing regime: neurons fire as soon as they leave their resting potential, thus displaying very low values of membrane potential when calculated (and sampled) over time.

Interestingly these pictures show that, up to the decisive point of the crises, the continuous measures look very similar, thus hinting at an instantaneous finite-size fluctuation causing the whole network to explode. Also, it is noteworthy that the new “hierarchy” set by the cascade is conserved in the non-propagative regime, while the propagative regime seems to have an overall reset effect.

Also, we see from these graphs that there is a particular time window where the variance of the mean membrane potential is larger for the most inhibitory-connected neurons, in both RS and FS populations (although it appears clearer for RS ones here, because of the need to rescale the FS colorbar to have comparable results). This increase of variance, while still present, is weaker and on a smaller time window in the average over connectivities compared to the average over noise realizations. This suggest connectivity plays a role in the intensity of the effect, although it remains qualitatively similar. We found that this time window defines the period when the network can actually switch to propagation: the high variance corresponds to different times when various realizations “explode”, and thus defines a region of instability.

A central point to raise here is that what makes the difference between propagative and non-propagative scenarios is most likely *not* an *infinitesimal* instability defined from a macroscopic perspective, i.e, that is due to a positive eigenvalue of a Jacobian defined from a large scale representation (Mean-Field for example), otherwise the non-propagative behavior would simply not be observable (as, except for chaotic dynamics, we do not observe unstable trajectories in phase space). Indeed, what differs between the various simulations is either the noise, or the connectivity realization, which may, or may not, bring the system to a point of instability. The external Poissonian drive, with *finite-size* fluctuations is thus constitutive of the scenarios we observe.

To gain more insight into the diversity of dynamics across neuron groups, we turn our attention to a measure of alignment, or synchronisation, namely the Kuramoto order parameter *R*.

#### Kuramoto order parameter

The Kuramoto parameter represents a degree of alignment, a value of 0 meaning there is no alignment while a value of 1 meaning everyone is perfectly align. We show in Fig.11(a)-(b) the Kuramoto order parameter *R* defined for each group of excitatory (RS) and inhibitory (FS) neurons in time, averaged over noise realizations (top row), and standard deviation over realizations (bottom row), in propagative (a) and non-propagative (b) scenarios (network connectivity held fixed).

**Figure 11:**
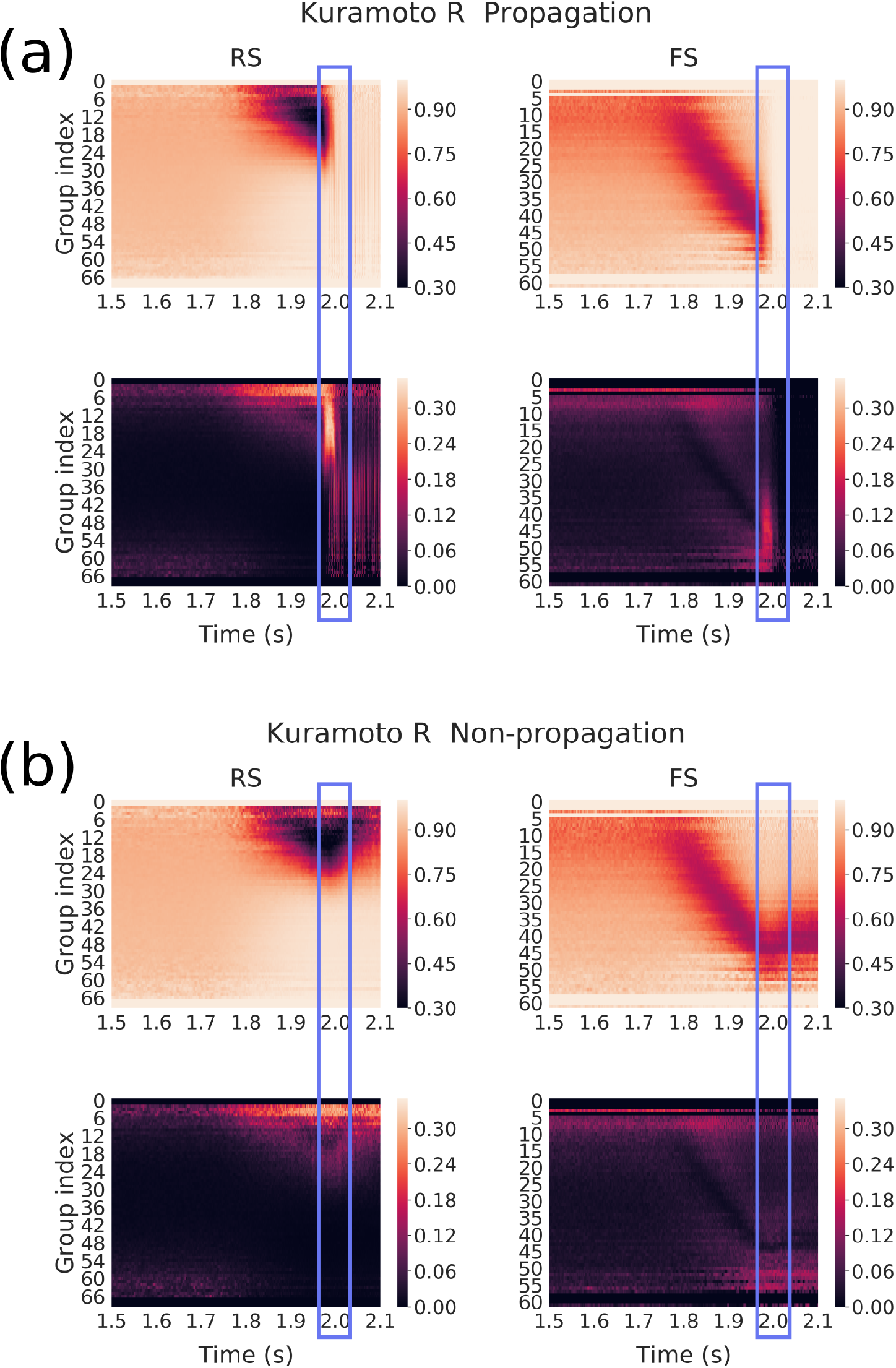
Kuramoto *R* of membrane potentials over subgroups of neurons (same network connectivity, different noise realizations) for each group defined as a function of their incoming inhibitory connections, averaged over 50 noise realizations (17 non-propagative and 33 propagative). Color maps correspond for each group to the average kuramoto parameter (top) and standard-deviation (bottom) across noise realizations in the propagative situations (a) and non-propagative situations (b) for both excitatory (RS) and inhibitory (FS) populations. The blue rectangle highlight the (time) region where the system either switches to a propagative regime, or remains stable. [FIGURE MODIFIED]

Again, we reproduce the results with 50 network connectivities, for both propagative and non-propagative scenarios, see Fig.12(a)-(b). We clearly see that the results are qualitatively similar, although with seemingly higher contrast than Fig.11.

**Figure 12:**
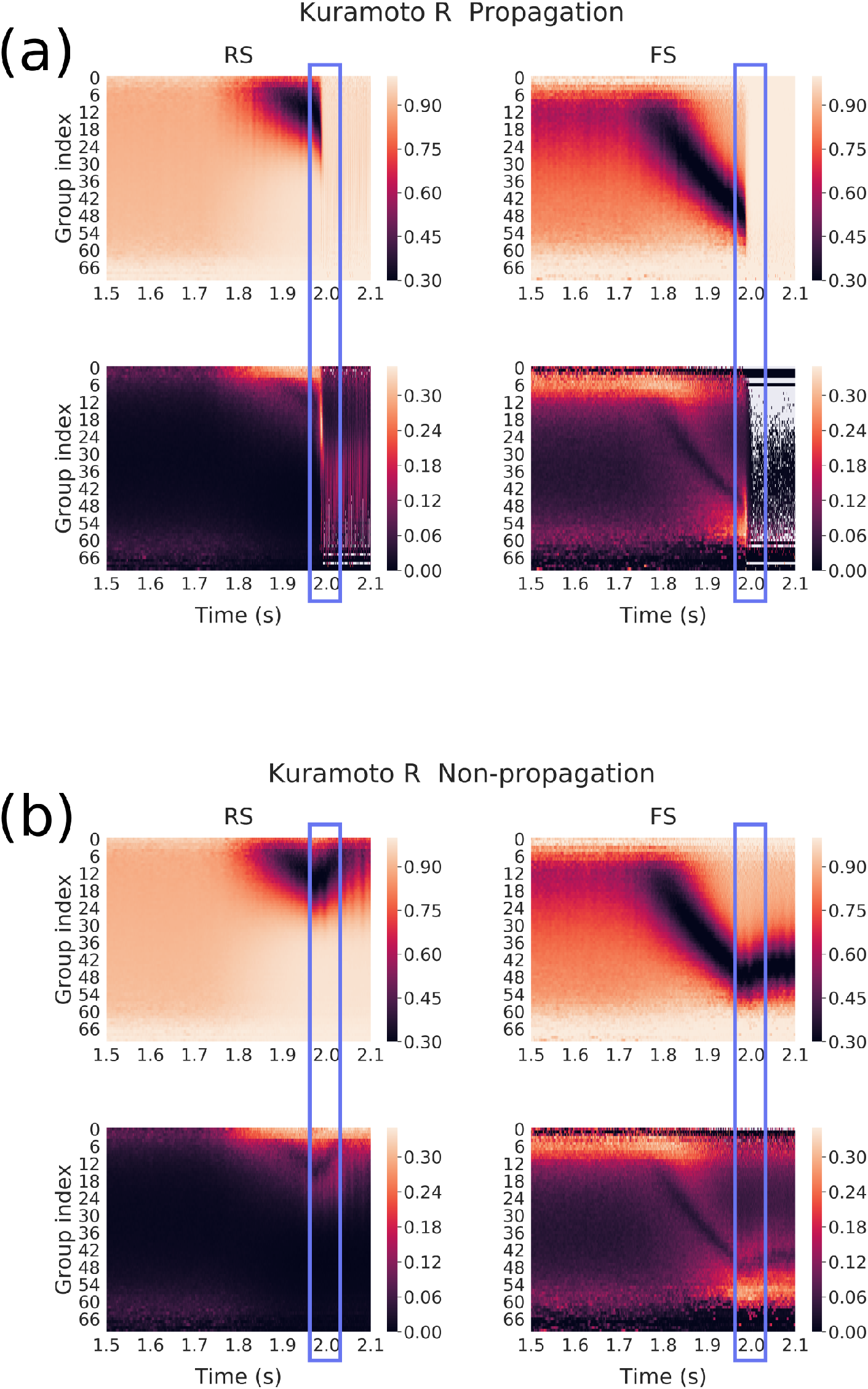
Kuramoto *R* of membrane potentials over subgroups of neurons (different connectivities) for each group defined as a function of their incoming inhibitory connections. Here, we averaged over 50 network connectivities for which we found a couple of noise realizations corresponding to propagative and non-propagative scenarios. Color maps correspond for each group to the average membrane potential (top) and standard-deviation (bottom) across different connectivities in the propagative situations (a) and non-propagative situations (b) for both excitatory (RS) and inhibitory (FS) populations. The blue rectangle highlight the (time) region where the system either switches to a propagative regime, or remains stable. [NEW FIGURE]

The cascade previously observed is clearly visible for the average *R*, in the form of a “desynchronization cascade”. For the propagative scenario, we note here a *recruitment* process between two radically different regimes having nonetheless alignment features: a fluctuation-driven asynchronous irregular (AI) dynamics, where membrane potentials are mostly conditioned by the balance of inhibitory versus excitatory inputs, and a crisis characterized by high spiking and membrane potentials clamped by refractoriness. Interestingly in the non-propagative scenario, it appears that the misalignment of the inhibitory neuron groups finally attained is fueled by the joint activity (of the network and the input), thus hinting at a out-of-equilibrium steady state (that continues until the end of the plateau of the perturbation, 1*s* later). From the standard deviation perspective, two main features are worth pointing. First, we again observe the instability window, characterized by high standard deviation between realizations in propagative scenarios. Secondly, we see that the two types of averaging leads to strikingly similar results, although slightly different quantitatively speaking, the average over connectivities leading to a higher contrast during the cascade. Therefore, our results are independent of both the noise realization and the specific connectivity, although an average over one or the other is useful to observe a typical case.

### 3.5 Dynamic versus static approach

We have seen that changing the slope and the amplitude of the signal alters the chances of triggering a crisis,thus hinting that the time evolution of the perturbation is central. Then we observed a hierarchical structure setting in from the point of view of continuous measures, following the perturbation. However, fundamental questions remain: how much of this latter phenomenon is actually dynamic? Would we find the same structures if we bombarded the network with a fixed input at, say, 80*Hz*? Can we observe the same dynamical structures for scenarios which are always, or never, propagative (no matter the noise or connectivity realization) ? This would indicate that the structures observed thus far might have little to do with the crisis phenomenology itself but would either be the mere results of strong conditioning of the network by the level of input (if static structures are similar), or simply not yield any explanation for the instability we observe (if always/never propagative scenarios show similar features).

We now turn our attention to Fig.13(a), which displays the static *µ*_*V*_ profiles in RS population obtained for fixed external inputs (“Stat.” curves), together with the profiles captured at the typical onset of the crisis, for various amplitudes: 60Hz (never propagative), 80Hz (sometimes propagative) and 100Hz (always propagative). The network realization is the same as previously analyzed, except when explicitly stated (Net. 2), where we refer to another connectivity realization. For the 80Hz scenarios with the first network (the one we have been investigating so far), we kept the splitting of the realizations between propagative and non-propagative, to highlight the potential differences of structures.

**Figure 13:**
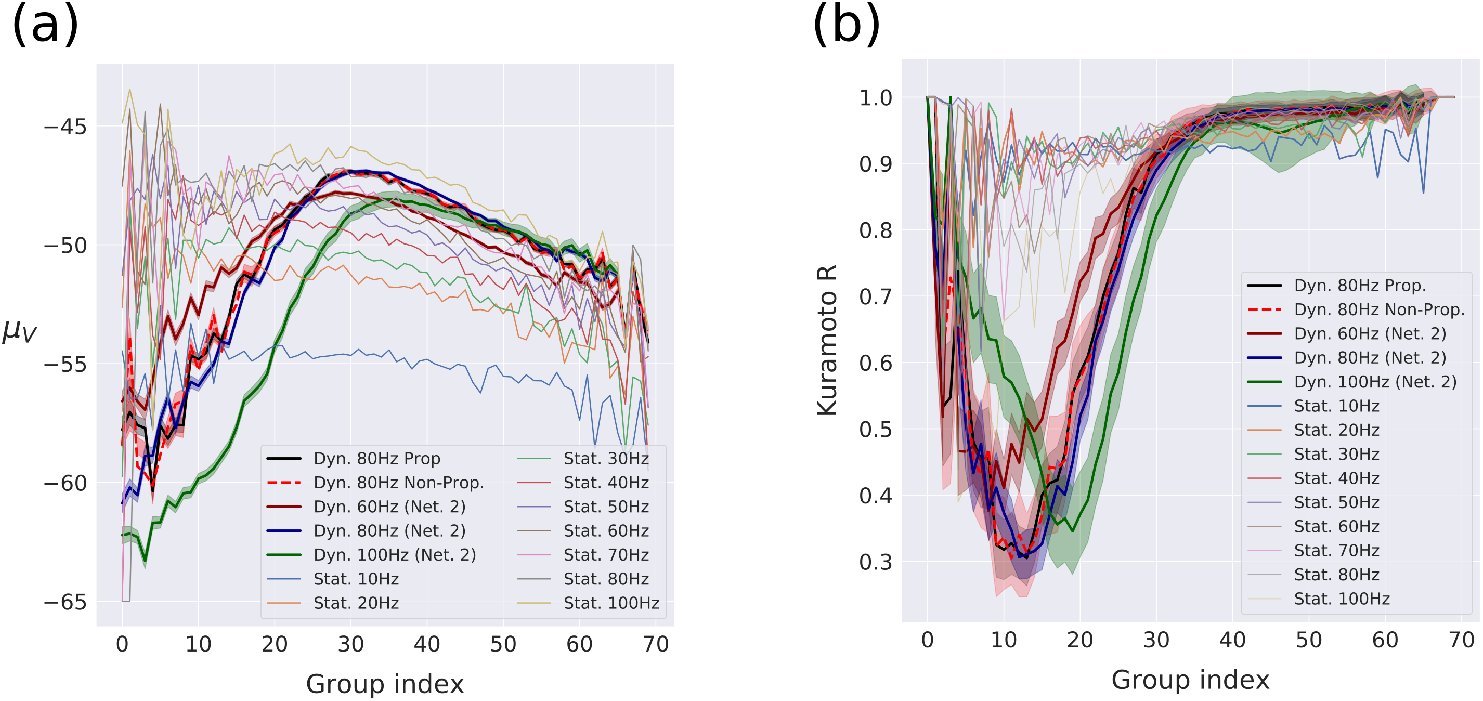
Steady-state and dynamical profiles of RS neurons for. **(a)** *µ*_*V*_ **and (b) Kuramoto** *R* over subgroups of neurons (same network connectivity (unless specified), different noise realizations), for fixed external input. The steady-states, called “Stat”, represent the stable activity without perturbation. They are drawn together with various profiles for different amplitudes of perturbation, called “Dyn”, captured right before typical time of crisis, at respectively 1950ms (60Hz), 1950ms (80Hz), 1930ms (100Hz, as the crises develops *before* 1950ms). Networks are the same as previously analyzed, except when stated Net. 2 which represent another network connectivity, for robustness. Standard errors estimated over noise realizations are shown in shaded areas.[FIGURE MODIFIED]

First, as previously observed, the profiles obtained for propagative versus non-propagative regimes are very similar for lower values of inhibitory connectivity.Then, we clearly see that the *µ*_*V*_ profiles extracted from the dynamical situations (hereafter called the dynamical profiles) are very different from the static ones.

Besides, it is worth pointing that the profile obtained for a 80Hz amplitude with a different random realization of the network (where all 50 noise realizations are put together, based on the previous observation that propagative and non-propagative scenarios show very similar structures) is very similar to those already shown, with small standard error, which, together with the previous observation that noise and network realizations seem to play similar roles, underlie a robust network phenomenology. Furthermore, we see that the profiles obtained for 60Hz, 80Hz and 100Hz amplitudes *are different*. The nature of their differences is of great interest for low indices, where we observe that 60Hz and 100Hz profiles are located on opposite sides of the central 80Hz profile: their ordering in this region is consistent with that of their response to the perturbation we have observed so far (see Fig.4). This said, the dynamical profiles yet show similar qualitative features : they all are non-monotonous and display two well-separated parts. Indeed, for low indices (until 30) *µ*_*V*_ is increasing with values starting around the lowest of the static profiles (10Hz), while their high indices part is more aligned with high static profiles. Interestingly, we see that for 60Hz and 80Hz the right part is well aligned with the static profile obtained for similar inputs. This does not seem to be the case for 100Hz, although the static input simulation displays some instability, which makes their comparison less relevant. Although it is not straightforward to link *µ*_*V*_ with the instantaneous regime, we have seen that low values can be associated with high firings (the neurons spending most of their time clamped at −65mV). This helps understanding what is happening here: for higher values of amplitude, the less inhibitory-connected neurons are firing more, and can thus entrain the rest of the network.

Fig.13(b) shows the Kuramoto order parameter aspect of the latter figure. Here the *R* profiles display structures quite different from those observed for *µ*_*V*_. Indeed, the various static profiles do not display such clear variability as for *µ*_*V*_, although little differences can still be observed: high inputs seem to show more variability in low indices, while ending at higher values for higher indices. More importantly, the dynamical profiles are here very different, from the static ones, and among themselves. Besides, propagative and non-propagative simulations show little differences here as well, and the profiles corresponding to same amplitude (80Hz) and different network architecture (Net. 2) also overlap here. Interestingly we can also observe that the 60Hz and 100Hz profiles are different and located apart from the 80Hz, although they also show different magnitudes of their inverted peaks. Given that the ordering of these magnitudes are not consistent with the various degrees of instability, we suggest that the *position* of the peak might be the most discriminating factor to establish whether the scenario is propagative. This would be consistent with the observations we made thus far, and confirm our previously suggested scenario: as the more we approach the center group, the more neurons are considered (Gaussian distribution), the green peak (100Hz) tells us that more neurons have undergone the desynchronization cascade we mentioned earlier, that is, more neurons have already “switched side” and entered a high firing regime, thus giving more inertia to the cascade phenomenon. The middle scenario (80Hz) would then sit on a *tipping point*, that is a point separating two radically different dynamical regimes of the system.

These latter observations show that, from the perspective of both mean membrane potential and Kuramoto order parameter calculated inside the groups formed from inhibitory in-degree, we are in the presence of a structured behavior which emerges from an intricate interaction between dynamics and architecture, and which cannot be recovered from static approaches.

### 3.6 Can seizure propagation be controlled by external inputs?

After having established that the structure of the dynamics allows or not the propagation of the paroxysmal perturbation, we now investigate whether we could use the previous finding of a strong instability window for the 80Hz dynamical scenario to alter the fate of the AdEx network dynamics. This approach is based on the following reasoning : we have observed, with a detailed analysis, that switching to one scenario or another is determined in a short time windows (just before the eventual crisis).Thus, we want to design a stimulation protocol to reduce the chance of crisis propagation, based on this observation, *but which does not require the same level of analysis*, hence making it applicable inline and without the need of extensive computational power. To do so, we will study the region around the crisis to determinate this relevant time window.

To achieve so, we apply a Gaussian stimulation, with 10 ms time constant, two different amplitudes (1Hz and 5Hz), positive or negative, *through a variation* of the external excitatory input (which depending on when the simulation is applied, can be the drive of 6Hz or the drive plus somewhere on the perturbation of 80Hz with a time constant of 100ms). For simulations performed under the same conditions, the stimulations were applied at different times as detailed in Tables 1(a)-(b). These tables show, for a total number of 100 simulations (with same network structure but different noise realizations), among which 72 were propagative, what relative percentage of simulations has undergone a triggering and a cancellation of crisis respectively.

**Table 1:** Triggered and prevented events: (a) Percentage of prevented events, from 72 initially propagative behaviors. Highlighted in orange ≥ 25% and in red ≥ 50%. The time of peak corresponds to the moment where the maximum of the stimulus is reached, the amplitude corresponds to a variation of the external input (see the main text) (b) Percentage of triggered propagation events, from an initial number of 38 non-propagative behaviors. Highlight in orange ≥ 25% and in red ≥ 50%.

|  |  |  |  |  |  |
| --- | --- | --- | --- | --- | --- |
| (a) | time of peak | +1Hz | +5Hz | -1Hz | -5Hz |
|  | t = 1500ms | 0.1806 | 0.1944 | 0.1528 | 0.1389 |
|  | t = 1850ms | 0.1389 | 0.1944 | 0.0972 | 0.1528 |
|  | t = 1950ms | 0.1528 | 0.2361 | 0.125 | 0.0694 |
|  | t = 1975ms | 0.0972 | 0.0 | 0.3472 | 0.3889 |
|  | t = 2000ms | 0.0139 | 0.0 | 0.25 | 0.5556 |
|  | t = 2500ms | 0.0 | 0.0 | 0.0 | 0.0972 |
| (b) | time of peak | +1Hz | +5Hz | -1Hz | -5Hz |
|  | t = 1500ms | 0.25 | 0.1786 | 0.2857 | 0.25 |
|  | t = 1850ms | 0.178 | 0.1786 | 0.2143 | 0.2143 |
|  | t = 1950ms | 0.0357 | 0.6071 | 0.2143 | 0.5 |
|  | t = 1975ms | 0.7143 | 1.0 | 0.25 | 0.28572 |
|  | t = 2000ms | 0.6071 | 1.0 | 0.0 | 0.0714 |
|  | t = 2500ms | 0.0 | 0.0 | 0.0 | 0.0 |

We see that it is possible to “reverse” the scenario from propagative to non-propagative in the time windows between 1975 ms and 2000 ms (and to vice versa, albeit for a larger time window) thanks to (or because of) the stimulation: as can be seen in (Table 1(a)) (see Table 1(b) for the opposite). A notably interesting case is that more than 50% of the crises are prevented if a stimulation of −5 Hz is applied in the same time window. This could open interesting leads in furthering qualitative comparisons between computational simulations and real-life situations, and eventually guide future interventions.

## 4 Discussion

In this computational work we studied the response of various spiking neural networks to paroxysmal inputs. We observed that the same networks can display various types of responses, depending on its nature (the neuron model used at its nodes), the shape of the perturbation (here we analysed particularly a plateau-like input with various slopes and amplitudes) and the realization of the random number generator. In the case of AdEx and CAdEx, two radically different responses to a qualitatively similar incoming excitatory perturbation are observed. Indeed, the latter could either recruit the excitatory population and thus allow the crisis to propagate to efferent areas, or be “controlled” by the activity of the inhibitory population, keeping the excitatory population at a low activity level, thus preventing further propagation. The response of the network depends not only on the amplitude of the perturbation but also on the “speed” at which it occurs. This is consistent with experimental observations (Saggio et al., 2020). Interestingly, in the case of a HH network, our investigations show very different network responses, where only the amplitude of the perturbation plays a role and where no variability on noise realizations was observed.

A rich literature shows that seizures can be classified according to their onset/offset features described by bifurcation types(Saggio et al., 2020; Saggio, Spiegler, Bernard, & Jirsa, 2017; Jirsa, Stacey, Quilichini, Ivanov, & Bernard, 2014). The most observed bifurcation at the onset of a seizure is a saddle-node bifurcation (Saggio et al., 2017), which is characterized by an abrupt change in the baseline of the electrophysiological signal (Jirsa et al., 2014). We observed in the current work that crisis are propagative in AdEx and CAdEx networks when they rise abruptly enough in the network. There is here an interesting correspondence revealing the importance of the onset of seizure dynamics, as it has been shown from a clinical point of view (Lagarde et al., 2018). It is worth noting that the absence of such phenomenology in HH networks (for the scenarios we considered) raises interesting questions in the modeling of seizure dynamics, but also more generally in neuronal networks : how the quantitative differences (number of variables) and qualitative differences (types of processes taken into account) in the single neuron models affect the global dynamics ? Are more precise models always the best in all respects ? This underlines the importance of the the choice (of model, of parameters): by modeling a neuronal network and observing a phenomenon which resembles reality, we are not testing whether the specific ingredients we chose *are* constitutive of this phenomenon, but *how they would be* if they were chosen a priori. It is only the systematic cross-model observations and comparisons that can yield such an answer as which are the necessary and sufficient ingredients to observe a given phenomenon.

Note that, in clinical observations, the most accessible measurements are made on a macroscopic scale. In the study proposed here, we observe the activities at a smaller descriptive scale by building a network of neuron models. We thus have a complex system of very high dimension, rendering a priori impossible to obtain a simple description of the dynamics, which motivates the statistical approach proposed here. With this type of analysis, we were able to track in time key features of the underlying dynamics, especially those supported by the structure of the network : inhibitory in-degree can be mobilized to explain global differences in network response. Indeed, we proposed a coarse-grained description of the network dynamics based on inhibitory in-degree, allowing us to capture internal processes that were not visible at first, and which play a significant role in the global out-of-equilibrium dynamics. We chose inhibitory in-degree as it was found to be the most influential aspect determining the firing rate (see Fig.5(a)). It is interesting to note inhibitory neurons were also the ones that had the highest firing rate (around 15*Hz*) while the excitatory neurons were way lower (around 2*Hz*) and the Poisson noise lower too (around 6*Hz* by construction). That difference could be the reason for the disparity in influence more than the nature of the neurons, and while it could be interesting to investigate, it is not in the scope of this study and does not change the main results as the categorization was only use as a tool to visualize the data.This opens the way to a flexible modeling framework of internal subpopulations, whose precision can be adapted to the most significant level of description, depending on the context and the questions asked. This is a first bottom-up step towards a coarser description of the system, and hence, may guide reliable modeling attempts at larger scales.

We have also established that not only this structure matters, but also its interaction with instantaneous finite-size fluctuations of the noise and the time evolution of the *global* dynamics. These are all constitutive of the observed behaviors, and none can be neglected to understand them.

Also, our results showed that, for the AdEx network, there exists a time window, characterized by a high variance across noise realizations, during which it is possible to reverse the behavior by applying an appropriate stimulation. The use of a stimulus to interrupt a seizure has been applied in the past in the case of absence seizure (Rajna & Lona, 1989). These results have been used as bases of computational studies at the scale of the EEG (Taylor et al., 2014). Computational work on the response of a network model to stimuli to disrupt seizure-like activities has shown the importance of the precise timing of the stimulation (Anderson, Kudela, Cho, Bergey, & Franaszczuk, 2007). Then, the use of electrode stimulation has been developed in rodents (Pais-Vieira et al., 2016). These different approaches have been implemented, including deep brain stimulation, vagus nerve stimulation (Boon, Cock, Mertens, & Trinka, 2018) and magnetic stimulation (Ye & Kaszuba, 2019). However, experimental recordings of the response to stimuli do not allow us to understand the mechanisms of large populations of neurons. Indeed, even if progress in calcium imaging or in multi-electrode arrays has made it possible since this last decade to record a large number of neurons simultaneously, we do not yet have access to the exact structure of the network they constitute. The study presented here is thus a proof of concept, based on a specific network model.

Finally, we also found that it is possible to “control” the propagation of the seizure by appropriate stimulation in a given time window. We think that this constitutes not only an important prediction of the model, but also a potentialimportant possibility of treatment of some types of intractable focal epilepsies. This prediction could be tested in future modeling work at the mesoscopic scale, with realistic connectivity between the focus and neighbring areas. Such a model could be used to test the hypothesis that appropriate stimulation in areas adjacent to the focus may prevent the propagation of the seizure.

Perhaps the most exciting perspective is that the same paradigm could be used experimentally to control seizures. This would require a system to detect the onset of the seizure in the focus, and another system to deliver appropriate stimuli in adjacent areas. Such a system could be applied to experimental models of focal seizures, to evaluate if such a paradigm could revert the propagation – and thus generalization – of the seizure. This could be another way of controlling seizures, not by suppressing the focus, but by making sure that the paroxysmal activity does not propagate.

## Acknowledgements / Funding information

This work was funded from the European Union’s Horizon 2020 Framework Programme for Research and Innovation under the Specific Grant Agreement No. 945539 (Human Brain Project SGA3) and the Centre National de la Recherche Scientifique (CNRS, France).

## Equations

**Adex model:**

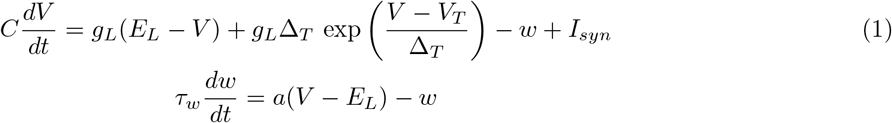

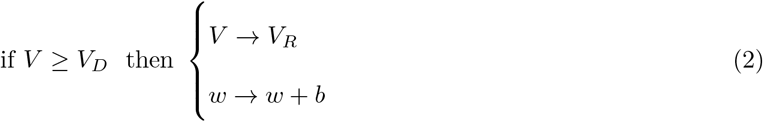

**CAdEx model:**

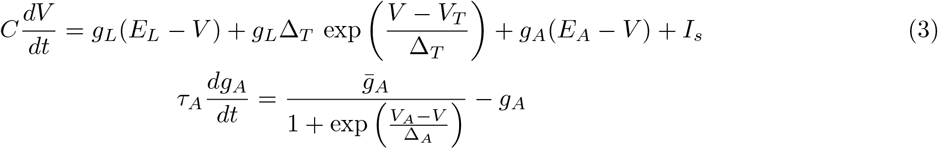

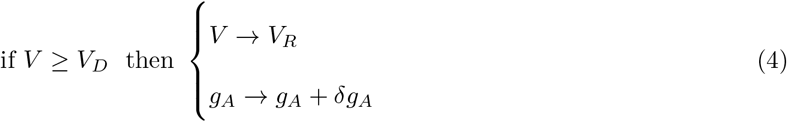

**HH model:**

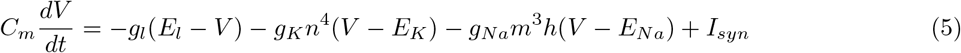

with gating variables (in ms):

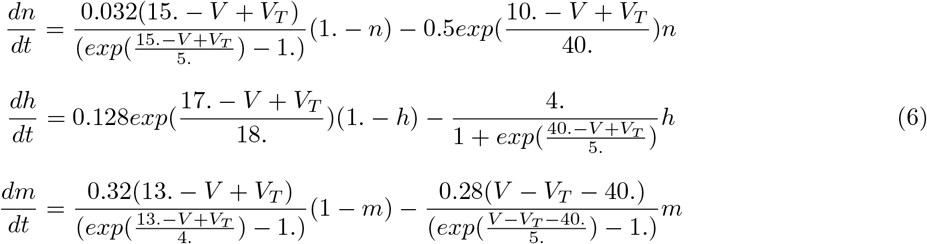

**Conductance-based synapses:**

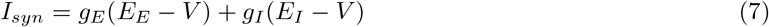

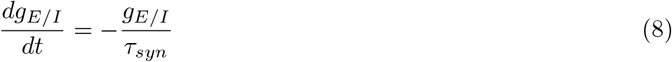

**External perturbation:**

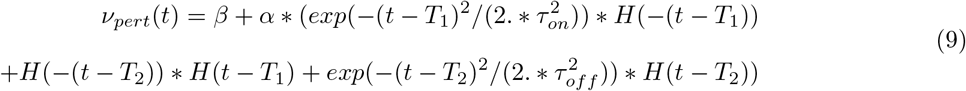

**Kuramoto order parameter:**

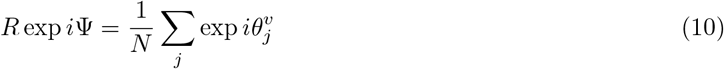

